# The GSS is an unreliable indicator of biological sciences postdoc population trends

**DOI:** 10.1101/171314

**Authors:** Christopher L. Pickett, Adriana Bankston, Gary S. McDowell

## Abstract

The postdoctoral research position is an essential step on the academic career track, and the biomedical research enterprise has become heavily dependent on postdoctoral scholars to conduct experimental research. Monitoring postdoc population trends is important for crafting and evaluating policies that affect this critical population. The tool most use for understanding the trends of the biological sciences postdoc population is the Survey of Graduate Students and Postdoctorates in Science and Engineering (GSS) administered by the National Center for Science and Engineering Statistics. To determine how well institutions tracked their postdocs, we analyzed the yearly changes in the biological sciences postdoc population at institutions surveyed by the GSS from 1980 to 2015. We find examples of large changes in the biological sciences postdoc population at one or a few institutions most years from 1980 to 2015. Most universities could not explain the data presented in the GSS, and for those that provided an explanation, the most common causes were improved institutional policies and more robust tracking of their postdocs. These large changes, unrelated to hiring or layoffs, sometimes masked population trends in the broader biological sciences postdoc population. We propose the adoption of a unified definition and titles for postdocs and the creation of an index to better assess biological sciences postdoc population trends.

**Abbreviations:**

NSF: National Science Foundation
NCSES: National Center for Science and Engineering Statistics
GSS: Survey of Graduate Students and Postdoctorates in Science and Engineering

## Introduction

Postdoctoral scholars are pivotal for the process of discovery in the biomedical research enterprise. Over the past several decades, the expansion of university research capacity and the availability of federal research grants have created a strong demand for highly skilled postdocs (Heggeness et al, 2016; Heggeness et al, 2017; National Institutes of Health 2016a, 2016b; Stephan, 2012). Despite this population growth and their importance to the research enterprise, comprehensive national-level data on the postdoc population is inaccurate or lacking. Within a single university, some postdocs may be categorized as employees whereas others are contractors, and some universities have multiple titles for postdocs reflecting differences in departmental practice, seniority and funding source (Ferguson et al, 2014). The variety of postdoc titles, while intending to clarify human resources policies or confer status on the postdoc, creates inconsistencies that introduce significant difficulties in collecting data on postdoc populations (Committee to Review the State of Postdoctoral Experience in Scientists and Engineers, 2014; McDowell, 2016). This is not a new problem: a 1969 report from the National Research Council states, “Although postdoctoral appointees were present on many campuses, their numbers and functions were not known nationally and, in many instances, were not even known to the host universities,” (Curtis, 1969).

That’s not to say the research community has not begun to tackle this problem. Spurred by National Academies reports and advocacy groups such as the National Postdoctoral Association, Future of Research and the Federation of American Societies for Experimental Biology, universities have created institutional definitions, established postdoc offices and crafted institutional policies to track their postdocs (National Academy of Sciences, 2000; nationalpostdoc.org; futureofresearch.org; http://www.sciencemag.org/careers/2002/02/faseb-adopts-definition-postdoctoral-fellows). However, these efforts are not uniform across the country: Definitions of who is a postdoc as well as titles, benefits, compensation and other aspects of the postdoc experience vary widely across U.S. institutions (Ferguson et al, 2014; Schaller et al, 2017).

At the national level, the National Science Foundation’s National Center for Science and Engineering Statistics (NCSES) conducts several surveys of the nation’s scientists and engineers to compile population data (http://www.nsf.gov/statistics/surveys.cfm), including the Survey of Graduate Students and Postdoctorates in Science and Engineering (GSS). While the GSS provides the most comprehensive assessment of trends in the postdoc population, it collects information on postdocs only from Ph.D. granting institutions and omits free-standing research centers and federal institutions that employ postdocs. The NCSES’ forthcoming Early Career Doctorates Survey may clear up some of these issues, and these data are expected in 2018.

While the GSS is generally considered the best tool available to measure the U.S. postdoc population, the GSS data collection practices are fluid. From 2007 to 2010, the GSS altered its methods of postdoc data collection (Einaudi et al., 2013). In 2014, the number of institutions surveyed in the GSS, known as the survey frame, increased so as to include more institutions with postdocs than had been evaluated before (Arbeit et al., 2016). These improvements enhance the accuracy of the survey and our understanding of the postdoc population, but they also introduce important caveats when evaluating postdoc population trends. Beyond this, the quality and consistency of the GSS postdoc data is dependent on the quality and consistency of postdoc information reported by individual universities. As a consequence, estimates of the entire U.S. postdoc population vary by 2-fold (Committee to Review the State of Postdoctoral Experience in Scientists and Engineers, 2014).

Here, we provide evidence that inconsistent reporting of institutional postdoc populations is a significant source of error in the GSS measurements. We focused on postdocs in the biological sciences as defined by the GSS (National Science Foundation, 2012). We highlight several institutions with widely fluctuating biological sciences postdoc populations that sometimes change by two-fold or more over a single year. We also found instances of dramatic changes in the postdoc population of one or a few institutions that mask contractions or expansions of the national biological sciences postdoc population. These data demonstrate the GSS biological sciences postdoc population data are unreliable when examining biological sciences postdoc population trends. The unreliability results from universities, funding agencies and advocacy groups failing to create a unified definition of a postdoc or put in place consistent policies to account for and track their postdocs. This has important consequences for those attempting to understand and change policies affecting postdocs. We call for a unified definition of a postdoc, consistent job titles and the creation of an index to more accurately follow trends in the biological sciences postdoc population.

## Results

When quantifying the postdoc population of an institution, a consistent set of policies on how to count postdocs is necessary to gain an accurate understanding of long-term changes in this population. An institution that changes its policies on who is considered a postdoc could have profound effects on the number of postdocs reported. To test this assertion, we examined the biological sciences postdoc populations at individual institutions as reported to the GSS between 1980 and 2015. We focused on the biological sciences postdocs reported in the GSS as this population has been the focus of recent influential reports on the role of postdocs in research (Committee to Review the State of Postdoctoral Experience in Scientists and Engineers, 2014; McDowell et al, 2014). Between 1980 and 2015, 379 of the 598 institutions surveyed by the GSS reported at least one biological sciences postdoc. To detect the largest changes in this population, we examined the 82 institutions that reported 100 or more biological sciences postdocs at least once between 1980 and 2015 (See Data collection and limitations). These 82 institutions represent 22 percent of all institutions reporting biological sciences postdocs, and they account for 77 to 86 percent of all biological sciences postdocs reported in the GSS over this time frame.

We identified large changes in an institution’s biological sciences postdoc population defined as a 2-fold increase or decrease in the postdoc population over the previous year. We identified 37 occurrences of an institution reporting 2-fold or more biological sciences postdocs than the prior year and 25 instances of an institution reporting 2-fold or fewer postdocs than the prior year (Fig. 1; Table S1). These 62 events happened across 27 institutions, or 7 percent of all institutions reporting a biological sciences postdoc from 1980 to 2015 (Table S1).

**Figure 1:**
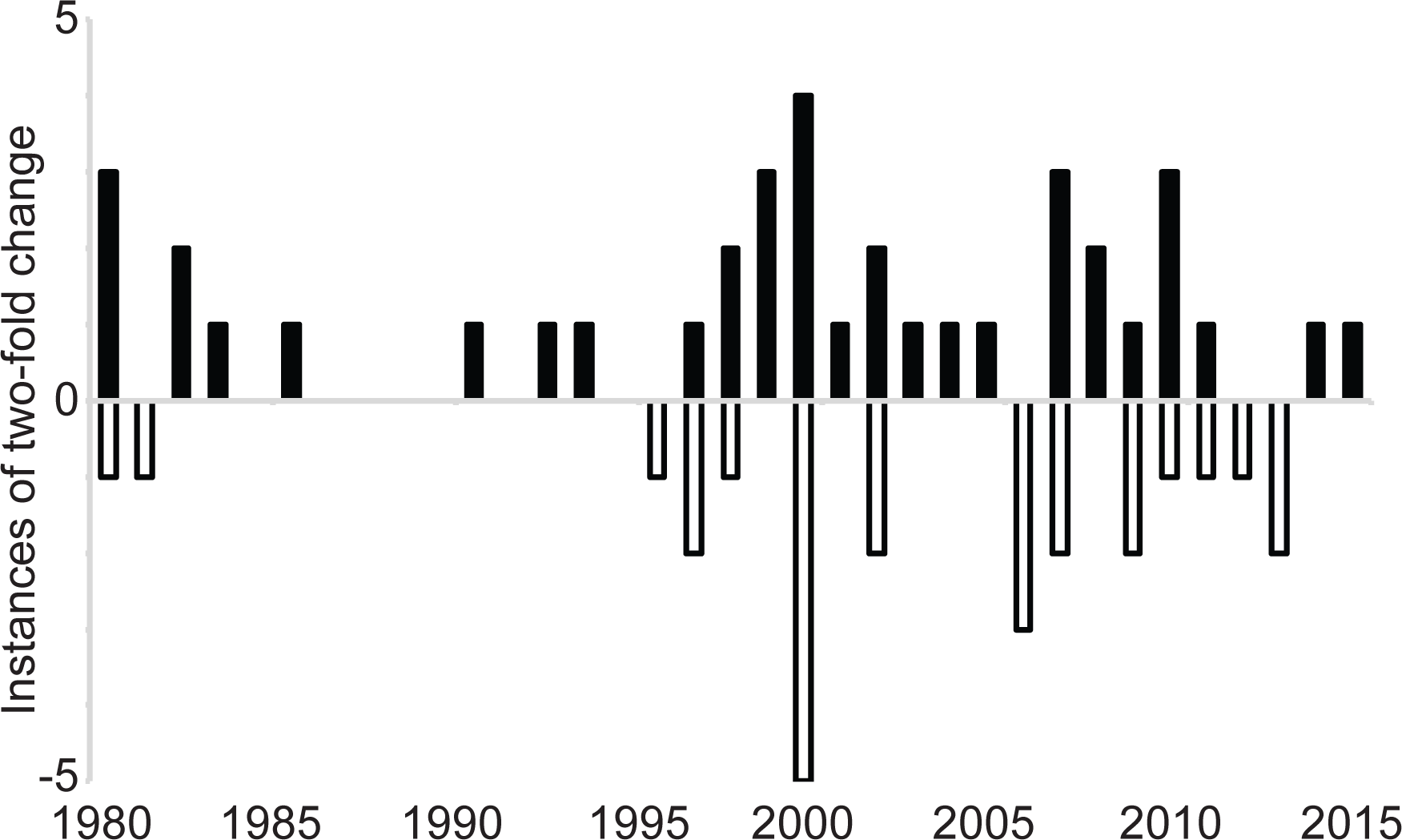
The number of 2-fold or more changes in the biological sciences postdoc populations of individual institutions reported in a given year between 1980 and 2015. Black bars indicate the number of instances of 2-fold increases and the white bars indicate 2-fold decreases.

Ten institutions had a single two-fold change in the biological sciences postdoc population between 1980 and 2015—eight reported a two-fold or more increase and two reported a two-fold or more decrease (Table S1). Of the institutions registering multiple twofold changes between 1980 and 2015, most are reciprocal changes (Table S1). For example, the University of Florida reported 108 biological sciences postdocs in 2000, 232 in 2001, and 106 in 2002 (Table S1). Similarly, the University of California, Riverside reported 92 biological sciences postdocs in 2009, 27 in 2010 and 89 in 2011 (Table S1). Other institutions followed these patterns, sometimes stretching the increase or decline across two or more years.

Of note, five institutions reported 0 biological sciences postdocs after reporting more than 75 at least two years before. Northwestern University (1997), the University of Maryland, Baltimore (2007) and the University of Tennessee Health Science Center (2009) each reported 0 biological sciences postdocs for a single year (Table S1). Brown University declined from 96 postdocs in 1999 to 0 in 2000, and it remained at 0 until 2005 when it increased to 88 postdocs (Table S1).

The GSS defines whether a postdoc is a biological sciences postdoc or a health fields postdoc based on the department or “organizational unit” they work in (National Science Foundation, 2012). Biological sciences organizational units are typically basic science departments while health fields units are more clinically focused. Because this definition is not necessarily reflective of the work done by the postdoc, one possible explanation for the 2-fold changes in the biological sciences postdoc population is that some postdocs were reclassified as health fields postdocs. To determine if reclassification as health fields postdocs could account for the observed changes, we analyzed the annual numbers of health fields postdocs at the institutions with 2-fold or more changes in their biological sciences populations. If biological sciences postdocs were reclassified as health fields postdocs, we would expect changes in the two populations to be of roughly equal magnitude but in opposite directions. Of the 62 2-fold changes identified in the biological science postdoc population between 1980 and 2015, the biological sciences and health fields postdoc populations changed in opposite directions 18 times (29%), but the changes were of approximately equal magnitude in only three of these instances (5%; Table 1). These data indicate that reclassifying biological sciences postdocs as health fields postdocs does not fully explain 95% of the 2-fold or more changes of institutions’ biological sciences postdoc population.

**Table 1:** Roughly reciprocal changes between biological sciences and health fields postdoc populations

| Institution | Postdoc population |  |  |  |  |  |
| --- | --- | --- | --- | --- | --- | --- |
|  | Year | Biological sciences<sup>a</sup> | Health fields<sup>b</sup> | Year | Biological sciences<sup>a</sup> | Health fields<sup>b</sup> |
| Augusta University/Georgia Health Sciences University | 1998 | 27 | 43 | 1999 | 69 (+42) | 3 (-40) |
| University of Florida | 2001 | 232 | 90 | 2002 | 106 (-126) | 188 (+98) |
| University of California, San Francisco | 2011 | 652 | 439 | 2012 | 325 (-327) | 689 (+254) |

<sup>a</sup>Number of biological sciences postdocs reported by the institution in the indicated year. Figures in parentheses indicate the change in postdoc numbers from the previous year.
<sup>b</sup>Number of health fields postdocs reported by the institution in the indicated year. Figures in parentheses indicate the change in postdoc numbers from the previous year.

Other institutional changes, however, could account for these changes. For example, institutions may have engaged in mass hiring or layoffs of biological sciences postdocs, and sometimes both within a few years of each other. Second, institutions may have changed their policies on how postdocs are classified and counted, resulting in dramatic changes in the reported biological sciences postdoc population. We contacted the 27 institutions reporting a 2-fold or more change in the biological sciences postdoc populations between 1980 and 2015 to assess whether these possibilities are responsible for our observation. Of the 62 2-fold or more changes, seven could be confirmed as being due to an institutional policy change (11%), and none were attributed to hiring or layoffs (Table S2; Supplemental Text). Twelve institutions, accounting for 23 2-fold or more changes (37%), could not explain the population changes in the GSS data as they occurred before the institution established reliable tracking techniques (Table S2; Supplemental Text). Eleven institutions, accounting for 27 2-fold or more changes (44%), did not recognize the postdoc population data from the GSS (Table S2; Fig. S1; Supplemental Text). This is could be due to differences in classification: institutions collect information on postdocs across all biomedical departments, while the GSS splits these postdocs into biological sciences postdocs and health fields postdocs. Three institutions, accounting for five observed changes (8%), did not respond to our request for information.

How likely are the 2-fold or more changes at individual institutions to affect the overall population trends of biological sciences postdocs? To answer this, we summed the change in the postdoc population due to the reported changes from schools that had a 2-fold or more change in their biological sciences postdocs over the previous year and compared it to the change in the national biological sciences postdoc population. For most years, the change in the postdoc population derived from schools with a 2-fold or more change accounts for only a small fraction of the change in the national population. However, 2004, 2005, 2006 and 2007 are exceptions. In 2004, the national biological sciences postdoc population increased by 91 postdocs, but a confirmed policy change at the University of California, San Francisco resulted in an increase of over 500 biological sciences postdocs over the previous year, a 4.8-fold increase (Table 2; Table S2). Thus, the policy change at UCSF masked a decline of 400 postdocs in the national biological sciences postdoc population. In 2005, the national biological sciences population increased by only 31 postdocs, but the Brown University biological postdoc population increased from 0 to 88, suggesting the change at Brown masks a 57-postdoc decline in the rest of the population (Table 2). In 2006, the overall population increased by 60, but three institutions reported a 2-fold or more change accounting for a decline of 194 postdocs (Table 2). This result indicates those institutions reporting a 2-fold or more change masked much larger growth in the national population (Table 2). In 2007, the increase in the number of postdocs accounted for in the institutions reporting a 2-fold or more change was nearly equal to the increase in the overall postdoc population, suggesting that postdoc population growth could have been flat (Table 2). These data points are important because the general sense was that the postdoc population increased across these years (Alberts et al., 2014; National Institutes of Health, 2012; Committee to Review the State of Postdoctoral Experience in Scientists and Engineers, 2014; Garrison et al., 2016). Yet it appears the expansion in the biological sciences postdoc population during this time may have been driven by reasons other than hiring.

**Table 2:** Dramatic changes in the biological sciences postdoc populations of just a few institutions can significantly alter the perceived trends of the population.

| Year | Institution(s) reporting 2-fold or more change | Combined change<sup>a</sup> | Change in national population<sup>b</sup> | Difference |
| --- | --- | --- | --- | --- |
| 2004 | University of California San Francisco | +509 | +91 | -418 |
| 2005 | Brown University | +88 | +31 | -57 |
| 2006 | Case Western Reserve University<br>Stony Brook University<br>University of Maryland Baltimore | -194 | +60 | 254 |
| 2007 | Case Western Reserve University<br>Stony Brook University<br>University of Maryland Baltimore<br>University of Massachusetts Medical School<br>The University of Tennessee Health Science Center | +287 | +302 | 15 |

<sup>a</sup>The sum of the changes at institutions that reported a 2-fold or more change in the biological sciences postdoc population over the previous year.
<sup>b</sup>The change in the national biological sciences postdoc population over the previous year.

Much of the individual institutional data reported in the GSS is likely accurate, but extremes in reporting from a few institutions make the total summed value for the national postdoc population unreliable. One way to evaluate the trends in the postdoc population without the complication of individual university policy changes would be to develop a postdoc population index that would analyze a select number of institutions to serve as proxies for population trends in the broader postdoc population, similar in concept to the Dow Jones Industrial Average or the S&P 500 indexes (Investopedia, 2015, 2017). This index would be defined, at least in part, by selecting institutions whose postdoc populations have changed within a reasonable percentage up or down over a specified time (Fig. S2).

As an example, we selected institutions that had a year-over-year change in the biological sciences population of less than 20 percent each year between 2001 and 2015 (Fig. S2). We chose 20 percent as a cutoff due to the current NCSES policy, instituted in 2010, that any university reporting a population change larger than 20 percent over the previous year is asked to ensure the information submitted is accurate (Einaudi et al., 2013). This indicates the NCSES views a 20 percent change as large enough year-over-year change to warrant closer inspection. We chose the time frame of 2001 to 2015 because the above data suggest the trends in the total postdoc population were strongly influenced by outlier events during these years (Table 2). We use 2010 as a useful demarcation in our analysis, as this is the year the national biological sciences postdoc population peaked and when important GSS data collection changes were fully implemented (Einaudi et al., 2013).

Only five institutions had year-over-year changes in their biological sciences populations that were less than 20 percent each year between 2001 and 2015, hereafter the 20/15 Index (20% threshold over 15 years). These five institutions accounted for between 8 and 10 percent of the national biological sciences postdoc population (Table 3). The trends in the biological sciences postdoc population of 20/15 Index institutions differed from the national biological sciences postdoc population in important ways. First, between 2001 and 2010, the 20/15 Index expanded in four years while it contracted in five (Fig. 2; Table 3). This is in contrast to the national biological sciences postdoc population which expanded for all nine years. Second, the longest period of growth in the 20/15 Index was two years (2006-2007), and the longest period of contraction was three years (2003-2005; Fig. 2; Table 3). Overall, the 20/15 Index increased 8.9 percent, whereas the total postdoc population increased by 27.6 percent over the same timeframe between 2001 and 2010 (Table 3).

**Table 3:** Characteristics of biological sciences postdoc population indexes relative to the national biological sciences postdoc population

|  |  | 20/15 Index<sup>a</sup> | 30/15 Index<sup>b</sup> | National |
| --- | --- | --- | --- | --- |
|  | Institutions included | 5 | 16 | 379 |
|  | Percent of national postdocs included | 8.3 - 10.0 | 19.2 - 22.7 | 100 |
| 2001-2010 | Years expanding<sup>c</sup> | 4 | 5 | 9 |
|  | Years contracting<sup>d</sup> | 5 | 4 | 0 |
|  | Longest consecutive increase | 2 | 2 | 9 |
|  | Longest consecutive decrease | 3 | 2 | 0 |
|  | Overall percent change<sup>e</sup> | 8.9 | 12.8 | 27.6 |
| 2011-2015 | Years expanding<sup>c</sup> | 1 | 0 | 1 |
|  | Years contracting<sup>d</sup> | 4 | 5 | 4 |
|  | Longest consecutive increase | 1 | 0 | 1 |
|  | Longest consecutive decrease | 4 | 5 | 4 |
|  | Overall percent change<sup>e</sup> | -6.4 | -9.2 | -8.5 |

<sup>a</sup>Institutions that changed 20 percent or less year-over-year each year between 2001 and 2015.
<sup>b</sup>Institutions that changed 30 percent or less year-over-year each year between 2001 and 2015.
<sup>c</sup>Number of years of population growth in the given time frame.
<sup>d</sup>Number of years of population decline in the given time frame.
<sup>e</sup>The biological sciences postdoc population in the indicated index in 2010 divided by the population in 2001 or the 2015 population divided by the 2011 population.

**Figure 2:**
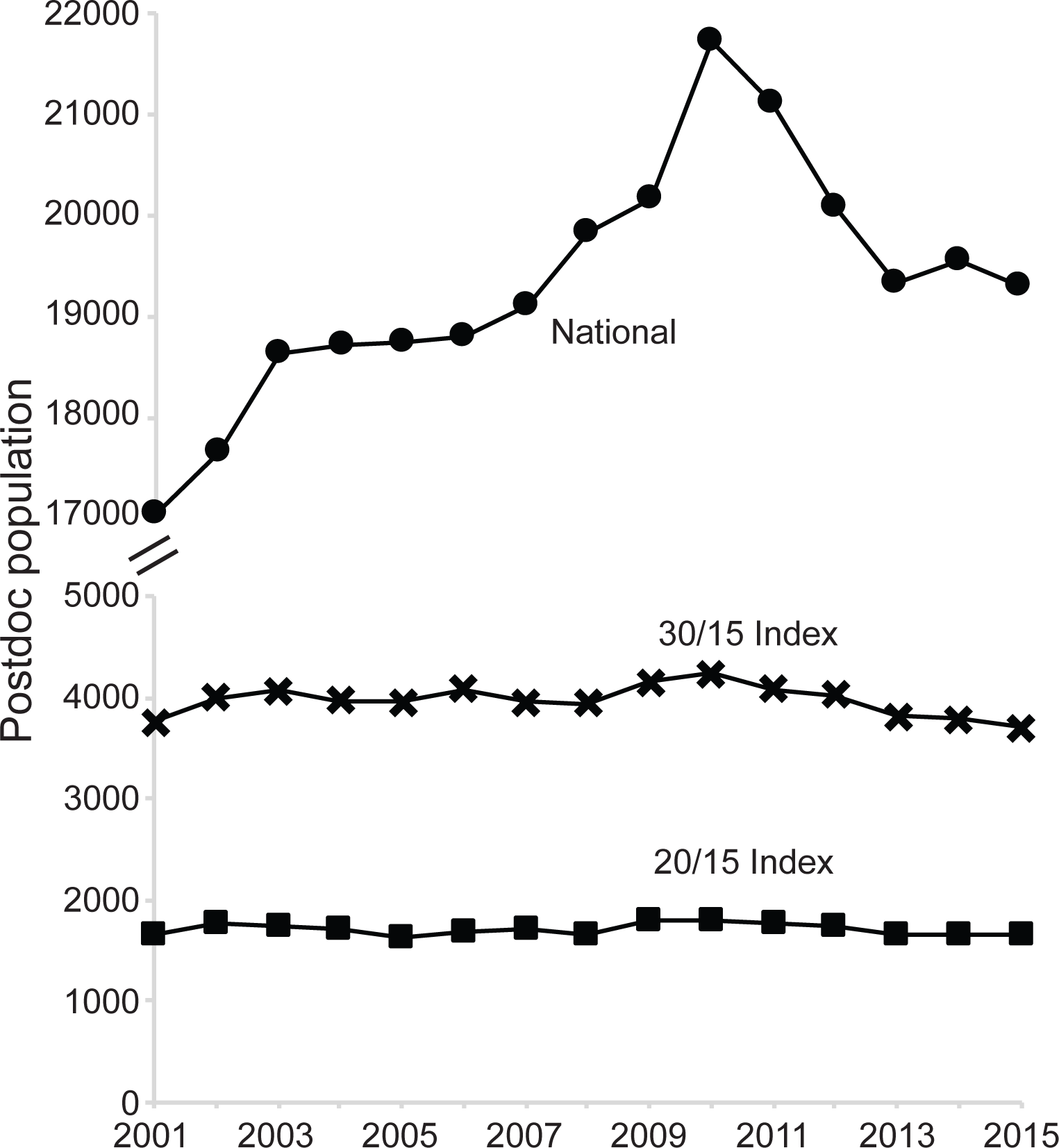
The population trends of the 20/15 Index (squares), 30/15 Index (Xs) and the national biological sciences postdoc population (circles) from 2001 to 2015. See text for parameters of each index.

Sixteen institutions were included if the threshold of year-over-year changes was increased to thirty percent over the same timeframe, hereafter the 30/15 Index (30% threshold over 15 years). These institutions accounted for 19 to 23 percent of the national biological sciences postdoc population. The 30/15 Index expanded in five years while it contracted in four (Fig. 2; Table 3). The longest period of growth in the 30/15 Index was two years (2002-2003 and 2009-2010), and the longest period of contraction was two years (2004-2005 and 2007-2008; Fig. 2; Table 3). Overall, the biological sciences postdoc population at the 30/15 Index institutions increased by 12.8 percent between 2001 and 2010 (Table 3). Thus, the 20/15 and 30/15 Indexes suggest modest growth of the biological sciences postdoc population between 2001 and 2010, with the population expanding for as many years as it was contracting. In contrast, the national biological sciences postdoc population, which includes institutions reporting more than 2-fold changes in their biological sciences postdoc populations, suggests a constant expansion in the population with an overall increase of more than 27 percent from 2001 to 2010.

The population trends in the 20/15 Index, the 30/15 Index and the total biological sciences population are largely the same from 2011 to 2015. All three measures declined for at least four of the five years between 2011 and 2015. Overall, the 20/15 Index and 30/15 Index declined by 6.4 and 9.2 percent respectively, while the national population declined by 8.5 percent between 2011 and 2015. These data are consistent with the observation that the biological sciences population declined broadly after 2010, and suggest the most recent updates to the GSS data collection methods have improved the survey (Einaudi et al., 2013; Garrison et al., 2016).

## Data collection and limitations

We queried the NCSES’s GSS through WebCASPAR (http://ncsesdata.nsf.gov/webcaspar/). We collected the number of postdocs at all institutions in all Broad Academic Disciplines from 1980 to 2015.

Data sorting and analysis was done in Microsoft Excel. First, we isolated the Biological Sciences values for each institution in the survey frame. We then used standard Excel formulas to identify institutions reporting 100 or more postdocs at least once between 1980 and 2015. Some institutional data was manually corrected: We combined the UMDNJ and Rutgers data because of their 2013 merger, the Georgia Health Sciences Center and Augusta University data because of their 2013 name change, and the Texas A&M University and Texas A&M Health Sciences Center data because of their 2014 merger. The City of Hope, Cold Spring Harbor Laboratory, Icahn School of Medicine, the Mayo Clinic Graduate School, Sanford-Burnham Medical Research Institute and The Scripps Research Institute were included in the GSS survey frame for only a subset of the 35 years between 1980 and 2015, and they were censored from our analysis.

To determine fold-change, the biological sciences postdoc population reported by an institution in a given year was divided by the biological sciences postdoc population in the year immediately prior for all values reported between 1980 and 2015 for the dataset described above. Values ≥ 2.0 indicated a 2-fold or more increase and values ≤ 0.50 were a 2-fold or more decrease of the postdoc population over a one-year period. Postdoc populations that decreased to 0 were classified as a >2-fold reduction and any increase from a 0 value was classified as a >2-fold increase. For identifying institutions for proposed indexes, we analyzed institutions with values for year-over-year change in their postdoc population between 0.833 and 1.2 for each year between 2001 and 2015 for the 20/15 Index and between 0.769 and 1.3 for each year between 2001 and 2015 for the 30/15 Index.

When reaching out to institutions to inquire about changes in their biological sciences postdoc population, our first point of contact was the institution’s postdoc office. These offices often answered our queries or referred us to other administrators with knowledge of the institution’s postdoc population. Offices were contacted at least three times to solicit a response. 24 of 27 institutions responded to our request for information on the data reported in the GSS.

## Discussion

Postdocs are a vital part of the research enterprise, and understanding the dynamics of this population is essential to developing policies that promote a vibrant research enterprise. Despite this, data on the postdoc population are unreliable. When considering the NSF surveys that gather data on postdocs, the authors of the 2014 Postdoc Experience Revisited report remarked, “[The committee] has little confidence in the accuracy of the absolute number of postdoctoral researchers, and it is particularly dubious about the quality of the information about postdoctoral researchers who are temporary residents and earned their Ph.D.’s in other countries. Nevertheless, the committee considers the available data to be a reliable indicator of trends over time. The gaps and flaws that exist are the same gaps and flaws that have existed for decades, so at least it may be supposed that the data possess some internal consistency,” (Committee to Review the State of Postdoctoral Experience in Scientists and Engineers et al., 2014).

Our findings challenge the assertion that the Survey of Graduate Students and Postdoctorates in Science and Engineering is internally consistent with regard to the biological sciences postdoc population and “a reliable indicator of trends over time”. Dramatic changes in the biological sciences postdoc population due to institutional policy changes have demonstrably distorted the trends of this population in the mid-2000s. Rather than a continuous increase from 1980 to 2010, our data suggest large changes at a few institutions in specific years gave the appearance of growth in the postdoc population during this time, but these changes may have masked population declines.

Concerns over the quality of data on the postdoc population are not new. Reports since the mid-1990s recommended improving data collection on postdocs (National Academy of Sciences, 2000; National Research Council, 1994,1998, 2005). We note that 50 of 62 observed 2-fold increases or decreases in an institution’s postdoc population occurred after 1995, possibly reflecting attempts by institutions to follow these recommendations and better track their postdocs. In addition to these recommendations, focused advocacy on behalf of postdocs by organizations like the National Postdoctoral Association began in the early 2000s (Sreenivasan, 2003). The recommendations made by these reports and organizations may be partly responsible for improvements in postdoc tracking.

We commend the institutions that have changed policies to better track their postdocs. These improvements in data collection are essential to better understand this population that is so critical to the biomedical research enterprise. However, as long as institutions individually make policy changes with regard to how they count and report their postdocs, the GSS will remain an imperfect source of information on the trends of the national postdoc population.

Further complicating this issue are policy changes and data collection updates made to the GSS itself. For example, a series of improvements in postdoc classification and accounting was implemented from 2007 to 2010, and the number of universities and institutions surveyed by the GSS expanded in 2014. These improvements in the GSS enhance the accuracy of the survey, but any comparison of trends in the postdoc population before and after these updates were implemented is suspect and should be treated with caution as they sample different proportions of the total postdoc population.

Collecting data on the postdoc population is difficult, but other groups have taken different approaches to examine this population. A study from the National Institutes of Health suggests there are roughly 30,000 postdocs funded by NIH grants (Pool et al, 2016). This is a broader accounting of the number of postdocs in the research enterprise relative to the GSS, mainly because this study is not confined to a defined set of universities; however, this study still misses some postdocs as it does not account for postdocs supported by organizations other than the NIH. The University of Michigan’s Institute for Research on Innovation and Science (IRIS) combines university human resources information with federal grant and census information to identify individuals working on research projects at universities (Zolas et al, 2015). This analysis has the potential to identify nearly all postdocs regardless of funding source, but to date, a limited number of universities have provided data to IRIS. One action all institutions could take to dramatically improve data collection on the postdoc population would be adopting simplified, unified Human Resources job classifications and a common postdoc definition and set of titles (Committee to Review the State of Postdoctoral Experience in Scientists and Engineers, 2014; Ferguson et al, 2014; Schaller et al, 2017). This would ensure each institution is consistently tallying and reporting data on the same population.

We propose the creation of an index of institutions that could provide a representative sample of the biological sciences postdoc population. The purpose of such an index is to understand the trends in the postdoc population by examining institutions with consistent postdoc definitions and policies for a prolonged period of time. The assumption of this index is that excluding institutions with high variability in their postdoc numbers will yield a clearer understanding of national postdoc population trends. We initially chose institutions whose postdoc populations changed less than 20 or 30 percent each year between 2001 and 2015—the 20/15 Index and the 30/15 Index, respectively. These indexes indicate that the postdoc population increased modestly from 2001 to 2010, with nearly the same number of years of expansion and contraction. This result is in contrast to the national biological sciences population, which increased every year during this time.

Analyzing postdoc population trends via indexes should be used in conjunction with analysis of the national postdoc population to identify discrepancies in population trends and highlight years of high variability in reporting. Presumably, as more institutions implement consistent practices for assessing their postdoc populations, more will be included in the indexes and index values will approach those of the overall postdoc population. For example, the 20/15 Index includes five institutions, but a 20/10 Index from 2006 to 2015 would include 13 institutions, and 33 institutions would be included in a 20/5 Index spanning 2011 to 2015 (Fig. S2). This could indicate that institutions are becoming more consistent with assessing their postdoc population.

Drawing conclusions from continuous datasets, such as the surveys conducted by the NCSES, should be done with caution. We propose the introduction of an index that is sensitive to the trends in the biological sciences postdoc population while eliminating artefacts introduced by changes not related to hiring or layoffs. This is not an ideal solution, but it is a workable solution until a unified definition and set of titles for postdocs are adopted by all U.S. institutions. Beyond improved postdoc data reporting, transparency in career outcomes and a greater inclusion of junior scientists in policy discussions are required to better understand the factors affecting the Ph.D. and postdoc populations (Dolan et al., 2016; McDowell et al., 2014; Pickett et al., 2015; Polka et al., 2015).

## Acknowledgements

We thank Shirley Tilghman, Melissa Vaught, Kenny Gibbs, Howard Garrison and Wally Schaeffer for helpful comments on an earlier version of this manuscript. C.L.P. and G.S.M. are supported by the Open Philanthropy Project and G.S.M. is supported at Manylabs by the Gordon and Betty Moore Foundation.

---

**Figure S1:** Graphs of the biological sciences postdoc population at individual institutions that reported a 2-fold change sometime between 1980 to 2015. The number of postdocs reported by an institution is on the y-axis and the year is on the x-axis. A blue line indicates a 2-fold or more increase in the postdoc population, a yellow line indicates a 2-fold or more decrease, and a black line indicates change was less than 2-fold.

**Figure S2:** The number of institutions included in an index varies based on the year-over-year threshold and the length of time analyzed. The x-axis indicates the percentile cutoff and the y-axis is the number of institutions achieving the cutoff. The endpoint of all timeframes is 2015 so the 5y line indicates the number of institutions achieving the indicated cutoff for each year from 2011 to 2015, 1Oy is between 2006 and 2015, 15y is between 2001 and 2015, etc. Red outlined black circles indicate the 20/15 and 30/15 Indexes.

